# Using readmers and hapmers in assessing phase switching after read error correction of Oxford Nanopore Sequences

**DOI:** 10.1101/2024.10.18.619002

**Authors:** Jean P. Elbers, David Horner, Tamara Löwenstern, Franco Laccone

## Abstract

Methods for sequence error correction can improve sequence accuracy; however, there can be unintended errors added during error correction. One such example is phase switching, whereby sequences derived from genomes containing more than one parental copy have contributions from more than one parental haplotype. Such switches are mistakes that can confound downstream analyses especially *de novo* genome assembly. While DNA sequences do not possess words such as linguistic languages, one can partition a DNA sequence into word-like objects called k-mers. K-mers include pieces of DNA sequence of k length that can describe various properties of DNA sequences. With regard to phase switching, there are k-mers present in one parental haplotype not found in other(s). These so-called hapmers can represent inaccuracies in that parental haplotype’s assembly but also correct, unique DNA variation. Here we investigated the effect of DNA sequence error correction on phase switching at the sequence/read level. Using several error-correction methods, we find all methods tested are similar to raw, presumably, phase-switch-free Oxford Nanopore Technologies (ONT) sequences in the percentage of readmers (k-mers from the ONT sequences) matching one parental haplotype’s hapmers. This work demonstrates an efficient method to assess if an error-correction method has introduced phase switching implemented in the Julia programming language.

## 1 Introduction

The advancement of sequencing technologies, such as Oxford Nanopore Technologies (ONT) sequencing, has revolutionized genomic research by enabling rapid and cost-effective long-read DNA sequencing. However, ONT sequencing reads have a tendency to have insertion and deletion (indels) errors, especially when the template sequence has homopolymers (Delahaye and Nicolas 2021; Huang, Liu, and Shih 2021). Indel sequencing errors can be corrected by a variety of computational methods, but it is not always clear whether non-intentional errors are introduced during the indel error-correction process as no method is perfect (Huang, Liu, and Shih 2021; Wang et al. 2020). One such type of unintentionally-introduced error is phase switching, a phenomenon where reads derived from multiple parental genome copies contain a mixture of more than one parental copy’s DNA nucleotides (Rhie et al. 2020). Phase switching may not be detrimental to all analyses, but it can confound downstream applications such as *de novo* genome assembly, whereby there could be loss of variation such as SNPs for phasing a genomic region resulting in that region being unphaseable (Konstantinos Kyriakidis, *pers. comm*.).

While DNA sequences do not possess words such as linguistic languages, one can partition a DNA sequence into word-like objects called k-mers (Waterman 1988). K-mers include pieces of DNA sequence of k length that can describe various properties of DNA sequences. With regard to phase switching, there are k-mers present in one parental haplotype not found in other(s). These so-called hapmers can represent inaccuracies in that parental haplotype’s assembly but also correct, unique DNA variation (Heng Li, *pers. comm*.). Further, during phase switching, there is a mixture of hapmers from heterozygous genomics regions that do not coexist on the same homologous chromosome.

While there are computational methods for estimating phase switching at the assembled contig or chromosome level (Li 2024; Rhie et al. 2020), here we investigate the effect of DNA sequence error correction on phase switching at the sequence/read level by comparing the k-mers in ONT reads (readmers) and percentage of hapmers from one or the other parental haplotype with human cell-line DNA. Using a computational approach implemented in the Julia programming language, we present a systematic evaluation of several error-correction methods.

## 2 Materials Methods

### 2.1 Cell Culture

We cultured HG002/RM8391 cells (NIST, Coriell), following the manufacturer’s culturing and expanding guidelines.

### 2.2 DNA Extraction, Library Prep, Sequencing

We extracted DNA from the HG002-cultured cells using automated extraction with EZ1 Advanced XL Robot (Qiagen) and EZ1 DNA Tissue Kit (Qiagen) and then performed left-size selection using Short Read Eliminator (SRE) Kit (Circulomics) to enrich reads greater than 25,000 bases. We incubated the samples with the SRE-reagent in a ratio 1:1 for 1 hour at 50°C to increase DNA yields. Next, we used the ONT SQK-LSK114 kit for ONT library preparation and the ONT-protocol for “Ligation sequencing 30kb human gDNA” (Version: GDH_9174_v114, revD, 10.11.2022) starting from the step “DNA repair and end-prep”. Finally, we sequenced the libraries on two R10.4.1 ONT PromethION flowcells using an ONT P2 Solo device connected to an ONT GridION.

### 2.3 Read Processing

We basecalled the resulting pod5 squiggles (i.e., the current raw output files from ONT sequencing) with Dorado 0.6.0 (ONT 2024) using the dna_r10.4.1_e8.2_400bps_sup@v4.3.0 basecalling model to initially generate SUP, raw ONT reads.

### 2.4 Error correction

While we are well aware there are numerous raw ONT sequence error-correction tools, we focused on the following as these were used in a prior analysis and were assessed to see if they might negatively impact phase switching at the sequence/read level. During error correction, we made a Snakemake workflow (Mölder et al. 2021) for processing and also used BBMap/ BBTools’s scripts (Bushnell 2024) such as partition.sh version 39.01, shred.sh version 39.01, and filterbyname.sh version 39.06 for various operations such as subsetting the reads, ensuring reads > 1,000,000 bases were split for effective downstream processing by one of the error-correction tools, Peregrine 2021 (Chin 2022), and downstream comparison of the same reads, respectively.

#### 2.4.1 Herro

Stanojević et al. (2024) developed the deep-learning, haplotype-aware ONT error-correction tool Herro. We used Herro commit # 5e7b70f and the version 0.1 R10.4.1 read-correction model. We first used seqkit 2.5.1 (Shen, Sipos, and Zhao 2024) to obtain read identifiers and minimap2 version 2.26-r1175 (Li 2018) to perform read overlapping, running the Herro developers’ create_batched_alignments.sh script. We then ran Herro inference (i.e., error correction) using a batch size of 64 and a single Nvidia A100 40GB graphical processing unit.

#### 2.4.2 Brutal Rewrite

Brutal Rewrite is an error-correction method that uses k-mers for error correction (Marijon, Sphor, and Limaset 2020). We used Brutal Rewrite commit # ad87f92, the graph algorithm for de Bruijn graph (Pevzner 1989) based error correction, and a k-mer length of 19 on Herro-corrected reads.

#### 2.4.3 Peregrine 2021

Peregrine 2021 (Chin 2022) is genome assembler similar to the Peregrine genome assembler (Chin and Khalak 2019) with both assemblers using sparse-hierarchical minimizers. We used Peregrine 2021 version 0.4.13 performing one round of overlapped-based error correction using the Brutal Rewrite-corrected reads as input and 6, 56, and 80 as values for the reduction factor, k-mer size, and window size for Peregrine 2021 settings, respectively.

#### 2.4.4 DeChat

DeChat is an error-correction method that we used on corrected reads but is designed for raw ONT reads (Li et al. 2024). We used default settings for version 1.0.0, using the Peregrine 2021-corrected reads as input and retaining only the *recorrected*.*fa* read DeChat output.

### 2.5 Phase switching analyses

#### 2.5.1 Read alignment

We used minimap2 version 2.28-r1209 with the following options to map raw, Herro, Brutal Rewrite, Peregrine 2021, and DeChat reads against either the maternal or paternal haplotypes separately from *hg002v1*.*0*.*1*.*fasta* (T2T_Consortium 2024): output “X” and “=“ in cigar strings, no secondary reads, soft-clipping of supplementary alignments, base-level alignment, output SAM format, and the high-quality, long-read preset. We then used SAMtools 1.9 (Li et al. 2009) to retain only primary alignments and those with mapping qualities equal to 60 and converted those SAM alignments into pairwise-mapping format (PAF) alignments with minimap2′s paftools.js sam2paf subprogram. Next, we retained the longest, primary alignment from each read and excluded those reads that Herro split into parts for Herro, Brutal Rewrite, Peregrine 2021, and DeChat reads.

#### 2.5.2 K-mers and hapmers

We developed a Julia programming language script (Bezanson et al. 2017) that takes read alignment information from minimap2 PAF files, FASTA files, and FASTA-index files of the reads and extracts ONT read k-mers (hereafter readmers) and associated k-mers unique to each haplotype (hereafter hapmers). We required that for a read alignment to be used for readmer and hapmer analysis, that query start and query stop had to be identical for maternal and paternal alignments (i.e., the same region of the read aligned to both haplotypes). Next, we iterated over each alignment meeting our criteria to extract the readmers and noted the total hapmers and matching hapmers for both maternal and paternal haplotypes. We processed the resulting tables with a Julia programming language script, converting the number of matches to percentages. For each read alignment, we aggregated the difference in the percentages of matching hapmers by taking the absolute value of the difference between the maternal and paternal matching percentages. For example, if we had a read alignment with 75% maternal hapmer matches and 25% paternal hapmer matches, then we would have 50% as the resulting value for that read alignment and likewise 50% if percentages were flipped between haplotypes (i.e., we would not have −50%). Finally, we generated histograms with a bin width of 1 from 1% to 100% with a cut-off of 6,000 read alignments per bin on y-axis or no cut-off. For all analyses we used a k-mer length of 15.

#### 2.5.3 Alignment statistics

Using the same reads for hapmer analysis, we realigned these sequences to both *hg002v1*.*0*.*1*.*fasta* assembly haplotypes simultaneously using minimap2 version 2.28-r1209 with the following options: output “X” and “=“ in cigar strings, no secondary reads, soft-clipping of supplementary alignments, base-level alignment, output SAM format, and the high-quality, long-read preset back to an assembly preset (i.e., lr:hqae and not lr:hq). We then used the program Bam Error Stats Tools (“best”, commit #c1c69bb, (Liu et al. 2024)) to analyze errors in the aligned reads.

#### 3 Results

We generated 15,048,314 raw, super (i.e., SUP) accuracy reads that we error corrected with Herro, resulting in 4,578,144 reads of which 582,208 were split parts of reads and not analyzed for phase-switching analysis. Brutal Rewrite does not filter out reads, but there were 4,490,689 reads remaining after one round of overlapping with Peregrine 2021 of which 532,757 were split parts and not analyzed for phase-switching analysis. DeChat did not filter out any reads from Peregrine 2021. Table 1 shows results for the number of read alignments meeting criteria for each correction method, prior to requiring the same read identifier for all 5 datasets for downstream hapmer analysis.

**Table 1:** Number of read alignments and percent out of total reads after filtering for alignments with mapping quality 60 and the longest, primary alignments. Herro, Brutal Rewrite, Peregrine 2021, and DeChat read alignments were further filtered for any reads split by the Herro algorithm. Coverage is estimated based on a haploid genome size of 3.1 billion bases

|  | Total Reads | Maternal Alignments | Paternal Alignments | N50 | Coverage |
| --- | --- | --- | --- | --- | --- |
| Raw | 15,048,314 | 4,029,514<br>(26.7772%) | 3,962,042<br>(26.3288%) | 21,268 | 40.3x |
| Herro | 4,578,144 | 1,733,156<br>(37.8572%) | 1,705,413<br>(37.2512%) | 24,437 | 26.3x |
| Brutal Rewrite | 4,578,144 | 1,733,152<br>(37.8571%) | 1,705,385<br>(37.2506%) | 24,437 | 26.3x |
| Peregrine 2021 | 4,490,689 | 1,724,488<br>(38.4014%) | 1,697,130<br>(37.7922%) | 24,152 | 25.7x |
| DeChat | 4,490,689 | 1,724,727<br>(38.4067%) | 1,697,341<br>(37.7969%) | 24,153 | 25.7x |

In general, the percentage of read alignments passing our filtering criteria of mapping quality 60 and only the longest, primary alignment increased with each successive correction method, except there were lower percentages for Brutal Rewrite than Herro (Table 1).

There were about 1,145,743 reads alignments that had the same query start and query stop for maternal and paternal haplotypes and were present in the raw, Herro, Brutal Rewrite, Peregrine 2021, and DeChat datasets. This 5-way intersection allowed us to keep reads identical but change merely what correction if any was applied to examine readmers matching hapmers.

Looking at Figure 1, one can see that the majority of the read alignments had close to 100% hapmers from a single parental haplotype suggesting little to no phase switching. The Peregrine 2021 method introduced a few taller, lower-percentage peaks, which could indicate a slight increase in phase switching at the read level. That being said, this was only a tiny increase relative to the almost 1 million read alignments in the rest of the histogram (Figure 2).

**Figure 1:**
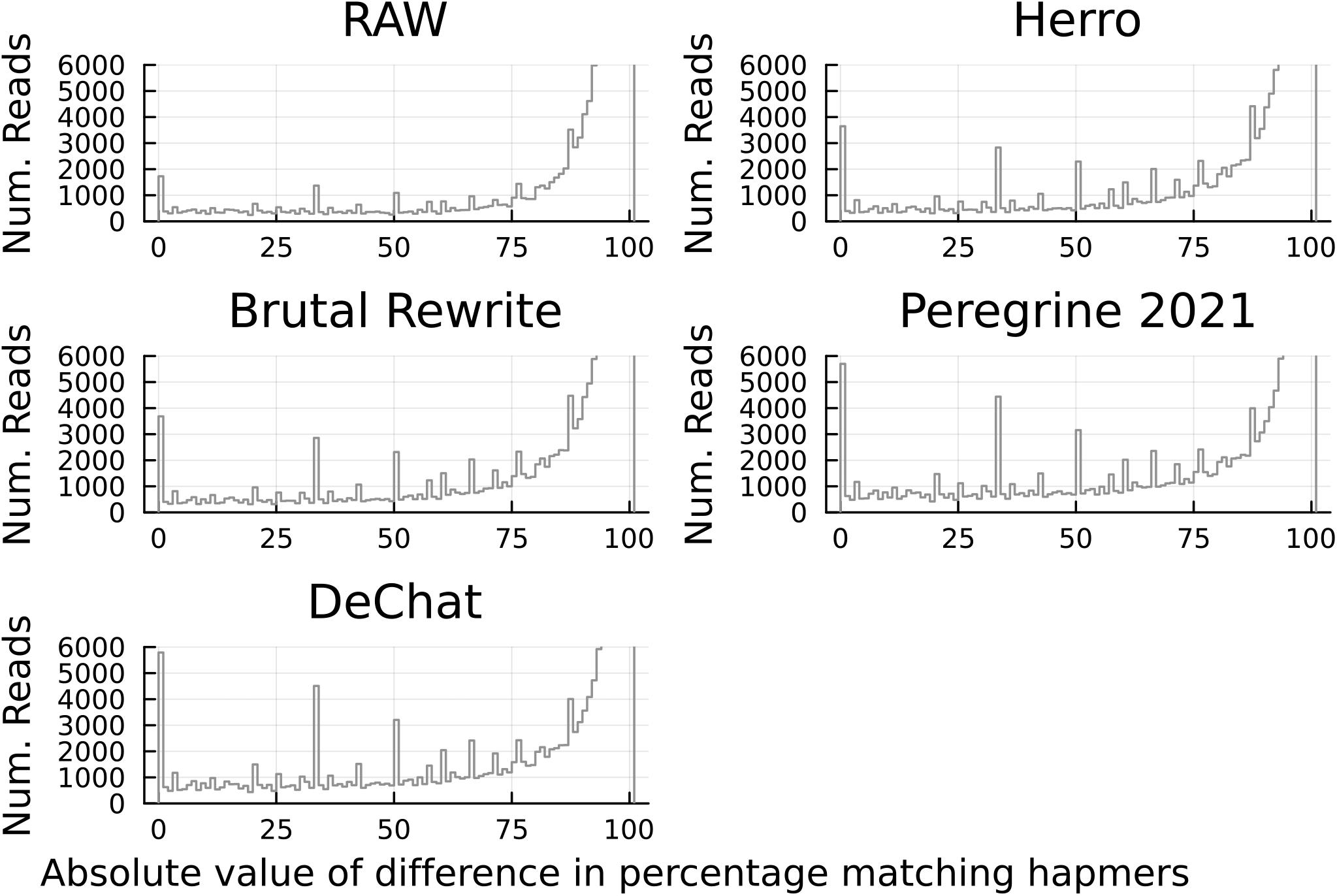
Histograms of absolute value of difference in matching hapmer percentage between maternal and paternal HG002 haplotypes. The y-axis was cut-off at 6,000 read alignments for visualization of low frequency data. The vast majority of occurrences were close to 100% suggesting that most of the read alignments possessed hapmers from a single haplotype and little to no phase switching.

**Figure 2:**
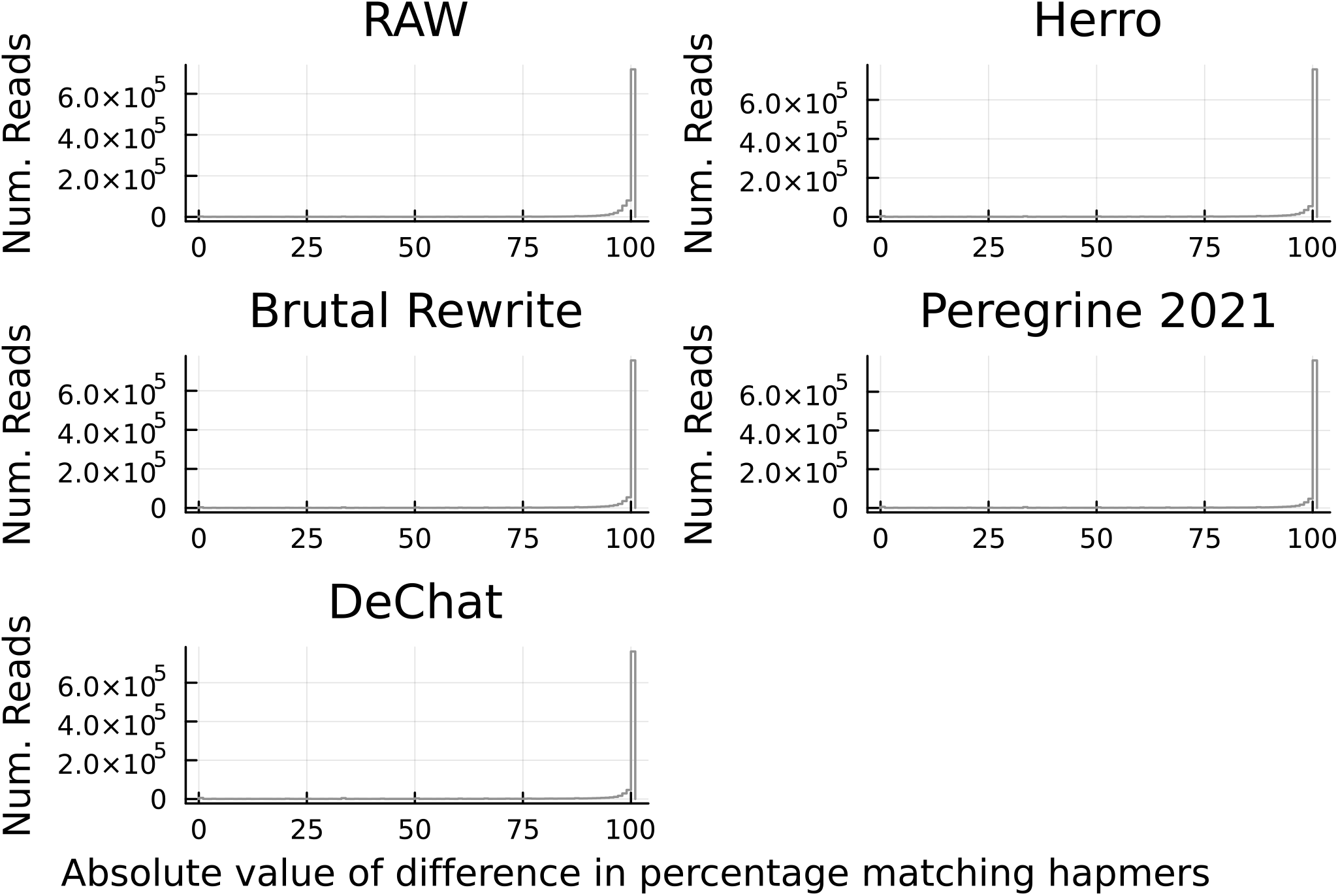
Histograms of absolute value of difference in matching hapmer percentage between maternal and paternal HG002 haplotypes. The y-axis is not cut-off and shows the staggering number of read alignments with almost 100% hapmers from a single haplotype.

Figure 2 shows in greater detail such a high proportion of the total read alignments coming from a single haplotype (i.e., how many read alignments had approximately 100% matching hapmers to a single haplotype).

There were some interesting scenarios whereby almost 12% analyzed, read alignments aligned in regions where 0 hapmers (i.e., homozygous genomic regions) were present (Table 2). Keeping in mind the distinction between a matching hapmer (i.e., a hapmer in a heterozygous genomic region matching a readmer in a read alignment) and a hapmer (i.e., a hapmer in a heterozygous region), we also observed read alignments with 0 matching hapmers but >0 total hapmers in the genomic region (Table 2).

**Table 2:**
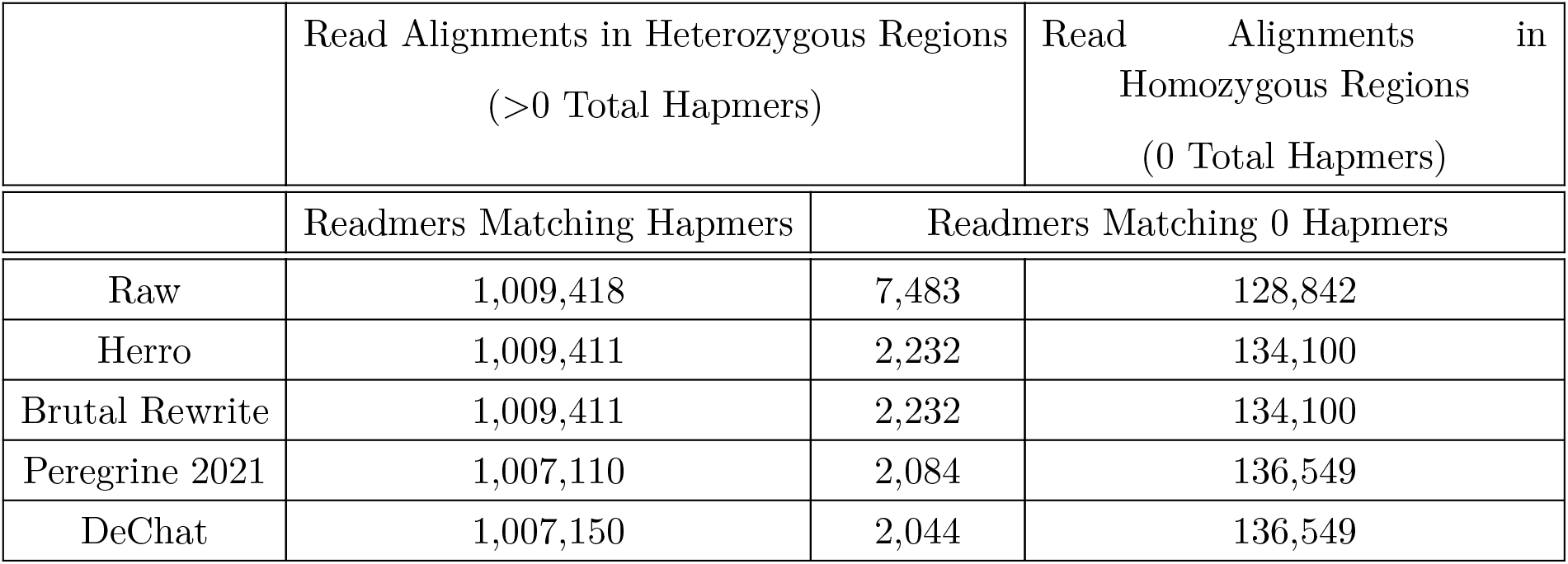
Number of read alignments (out of 1,145,743 total) based on whether the reads were aligned in heterozygous or homozygous regions.

**Table 3:** Alignment statistics from the program Best (Bam Error Stats Tools) of reads used in hapmer analyses. Abbreviations: Compres for compressed, Kbp for Kilobase pair, Inser for Insertion, Del for Deletion, Hp for homopolymer.

|  | Identity | Identity Quality Value | Gap Compres Identity |
| --- | --- | --- | --- |
| Raw | 0.980335 | 17.062956 | 0.984901 |
| Herro | 0.999637 | 34.396470 | 0.999770 |
| Brutal Rewrite | 0.999637 | 34.398948 | 0.999772 |
| Peregrine 2021 | 0.999726 | 35.619356 | 0.999830 |
| DeChat | 0.999741 | 35.862487 | 0.999842 |
|  | Matches per Kbp | Mismatches per Kbp | Non-hp Inser per Kbp |
| Raw | 985.021819 | 7.253553 | 2.826602 |
| Herro | 999.725438 | 0.052275 | 0.033748 |
| Brutal Rewrite | 999.725298 | 0.051433 | 0.033581 |
| Peregrine 2021 | 999.790755 | 0.031596 | 0.024639 |
| DeChat | 999.803664 | 0.027125 | 0.023584 |
|  | Non-hp Del per Kbp | Hp Inser per Kbp | Hp Del per Kbp |
| Raw | 3.413819 | 1.954716 | 4.310809 |
| Herro | 0.044347 | 0.055095 | 0.177940 |
| Brutal Rewrite | 0.044946 | 0.054916 | 0.178323 |
| Peregrine 2021 | 0.026350 | 0.040332 | 0.151300 |
| DeChat | 0.022643 | 0.039365 | 0.146568 |

Using the same reads used in the hapmer analysis but looking at the alignment statistics such as alignment identity, matches/mismatches per kilobase, and non-homopolymer/homopolymer insertion/deletions per kilobase: we saw that the raw reads had much lower identities, matches per kilobase, and higher values for other categories compared to all other corrected reads. Each successive correction method seemed to improve the alignment statistics, but one notable thing was that Brutal Rewrite had a lower non-homopolymer insertion rate per kilobase than Herro, but a higher non-homopolymer deletion rate per kilobase. The same difference between insertion and deletion rates occurred for homopolymers between Brutal Rewrite and Herro.

## 4 Discussion

### 4.1 General

The findings presented here provide significant insights into the effects of various error-correction methods on phase switching at the read level in ONT sequencing. One of the key observations is that all tested error-correction methods—Herro, Brutal Rewrite, Peregrine 2021, and DeChat— exhibited similar percentages of readmers matching one parental haplotype’s hapmers compared to raw ONT sequences. This suggests that while these methods improve overall sequence accuracy, they may not significantly exacerbate phase switching issues.

### 4.2 Phase Switching Insights

The results indicate that the majority of read alignments primarily matched hapmers from a single parental haplotype, which suggests minimal phase switching occurred. However, the presence of taller, lower-percentage peaks in the Peregrine 2021 reads (Figure 1) raises questions about Peregrine 2021′s haplotype awareness. Although this increase was minor, it highlights the need for further investigation into how certain error-correction tools can inadvertently introduce errors during the correction process. Peregrine 2021 is very fast in read overlapping as it avoids all-versus-all read overlapping (i.e., the computationally expensive process of overlapping each and every single read) through its unique, sparse-hierarchical minimizer index that helps find overlap candidates instead all-versus-all overlapping. Perhaps not doing all-versus-all overlapping increases increases phase-switching ever so slightly. We experimented with alternative values for the reduction factor and window size when running Peregrine 2021, we used 4 and 64, respectively, but this did not change the overall pattern observed in the initial Peregrine 2021 hapmer analysis. As Peregrine 2021 uses hand-coded heuristics in error correction, it may not have performed as well as Herro, which is a deep learning based method, especially given that deep learning methods outperform hand-coded heuristic methods on low coverage data used here (Jason Chin, *pers. comm*.).

Interestingly, the analysis revealed a subset of read alignments that had no matching hapmers to readmers in regions where hapmers existed. This could be indicative of sequencing errors from those genomic regions, misalignments, or perhaps even low-levels of contamination. While we did not further investigate these situations, understanding these discrepancies could help refine error-correction strategies.

### 4.3 Error Rates in Different Correction Methods

An intriguing finding is the observed difference in non-homopolymer insertion and deletion rates between the Brutal Rewrite and Herro methods. Specifically, Brutal Rewrite demonstrated a lower non-homopolymer insertion rate per kilobase but a higher non-homopolymer deletion rate compared to Herro. This raises questions about the underlying mechanics of these error-correction algorithms. It may suggest that Brutal Rewrite is more conservative in adding new bases but potentially more aggressive in removing bases that it deems erroneous. As our estimated coverage was only 26.3x for the reads used in Brutal Rewrite correction, perhaps higher coverage would have been beneficial. Further in regions of low-complexity repeats, Brutal

Rewrite tends to select the shortest repetitive sequence (Antoine Limasset and Pierre Marjion, *pers. comm*.).

Similar patterns were observed in terms of homopolymer regions, where the insertion and deletion rates differed between the two methods. These variations highlight the importance of understanding the specific behaviors of each correction tool, as they may influence downstream analyses, such as variant calling and *de novo* genome assembly.

### 4.4 Future Directions

Future studies should focus on further characterizing the mechanisms behind phase switching and error introduction across different correction methods. A more comprehensive understanding of how each tool interacts with specific genomic features—such as homozygosity and heterozygosity—could lead to improved error-correction algorithms that maintain haplotype integrity while enhancing sequence accuracy.

### 4.5 Conclusion

In conclusion, while current methods show promise in maintaining haplotype fidelity and reducing sequencing errors, ongoing research is essential to refine these tools and their applications in genomics. Addressing the discrepancies noted in this study will be crucial for advancing our understanding of error-correction for ONT sequences.

## 5 Acknowledgements

We would like to acknowledge the Medical University of Vienna’s High Performance Computing cluster for computing resources, Konstantinos Kyriakidis for thoughts regarding the effect of phase switching on *de novo* genome assembly, Heng Li for thoughts regarding hapmer and readmer error rates, Jason Chin for thoughts regarding Peregrine-2021, and Antoine Limasset and Pierre Marjion for thoughts regarding Brutal Rewrite.

## 6 Data Availabilty

Code documenting how analyses were conducted is available at https://github.com/jelber2/hapmers. Raw, basecalled SUP accuracy reads are available in unaligned BAM format at the following Zenodo repository https://doi.org/10.5281/zenodo.13841954.

## 7 Author contributions

JPE executed all analyses and wrote the manuscript. DH cultured HG002 cells. TL performed DNA extraction, library preparation, and sequencing. FL reviewed the manuscript.

